# Organization of the trilobite-type brain may have curtailed its survival

**DOI:** 10.64898/2026.03.24.713993

**Authors:** Nicholas J. Strausfeld, Xianguang Hou, Frank Hirth

## Abstract

Cambrian stem mandibulates and chelicerates have provided robust evidence of brains that conform to a cerebral ground pattern also defining extant euarthopods. But information about the brains of trilobites or their immediate relatives remain a mystery. Here we describe the detailed organization of the specialized brain of *Xandarella spectaculum*, a member of the subphylum Trilobitomorpha which went extinct at the end Permian. While retaining the pre-Cambrian cephalic ground pattern as three seriate domains, empirical evidence indicates clade-specific accentuation of some and loss of other neural arrangements typifying extant mandibulates. The trilobitomorph deutocerebral organization is dominated by exaggerated glomerular centers wherreas its prominent lateral eyes supply minute visual centers. Paired ocelli are associated with the forebrain’s prominent prosocerebral bridge, an essential center for path integration. Together these modifications indicate neural organization typifying some extant benthic crustaceans and even hexapods that engage in foraging behaviors. With reference to existing neurological attributes of living pancrustaceans major divergences of the trilobitomorph brain from the ancestral cerebral ground pattern may have weakened its resilience to geo-ecological disruptions and hence the final extinction of the clade to which trilobitomorphs belong.

**Significance:** Trilobites and their trilobitomorph relatives are iconic arthropods that appeared early in the Cambrian more than half a billion years ago. Why they went extinct 300 million years later when other arthropods did not has been a mystery not helped by the calcareous exoskeleton of trilobites generally excluding soft tissue preservation. However, the trilobite sister species *Cindarella* and *Xandarella* provide exquisite fossilized nervous system. Identifiable neuropils and even discrete connections show detailed cerebral organization. While the basic seriate patterning of today’s pancrustaceans and trilobitomorphs align, profound accentuation of certain centers and the major loss of others in the trilobitomorph brain show that it was already adapted for supporting grazing behaviors but was locked out of re-evolving adaptations that may have allowed survival of provincial and global extinctions.

## Relevance of Trilobitomorpha, their external morphology and optical organization

Cambrian fossils have provided robust evidence of brains that conform to a cerebral ground pattern also defining extant euarthopods**^1, 2^**. However the brains of trilobites or their relatives have been inaccessible, thus providing no insights at all as to why they finally went extinct at the end Permian**^3^**.

Since the first identification of fossilized brains in Cambrian euarthropods**^4–6^**, subsequent discoveries of these rarities have provided further evidence supporting genealogical correspondences between the brain organization of stem euarthropods and that of modern taxa**^1, 2, 7–9^**. Neuroanatomical diversifications of morphogenetically fixed cerebral domains, recognized in extant euarthropods and indicated in fossilized species, demonstrate that already in the lower Mid-Cambrian the distinctive cerebral arrangements of radiodontan, mandibulate, and chelicerate ground patterns correspond to, respectively, cerebral arrangements typifying modern Onychophora, Pancrustacea, and Chelicerata**^10^**. Such neuroanatomical correspondences provide important evidence for resolving the phylogenetic placement of fossils within the arthropod tree of life**^8, 9^**.

In contrast to Mandibulata and Chelicerata, a third major clade of Cambrian euarthropods did not reach the Holocene. This was Artiopoda, erected by Hou and Bergström in 1997 and diagnosed by them as “lamellipedian arthropods with trilobite-like appearance, broad tergum, and usually a complete set of fairly undifferentiated post-antennal appendages”**^11,12^**. Artiopoda includes Trilobitomorpha, exemplified by the iconic trilobites**^3^** whose biomineralized (calcified) exoskeletons generally impeded fossilization of soft tissues. This condition contrasts with that of numerous members of Trilobitomorpha that possessed uncalcified exoskeletons more amenable to occasional preservation of soft tissue during fossilization**^13^**. Based on the overall similarity of trilobitomorph shapes and overall appendicular uniformity, remains of neural tissue in one taxon could likely be representative of the clade as a whole. Here we describe neural traces in two non-biomineralized trilobitomorphs. One, *Xandarella spectaculum***^11,12^** provides exceptionally well-defined traces of a brain as well as prosocerebral ocelli and protocerebral lateral eyes. The other is *Cindarella eucalla*, its close relative**^13^**, which provides clearly defined segmental ganglia.

The initial 1991 denomination of *X. spectaculum***^11^** and its subsequent 1997 description in the context of the Chengjiang euarthropod fauna**^12^** leave little to be added. As described in earlier reports**^11, 12^**, the overall dorsal view of *X. spectaculum* shows a teardrop-shaped body comprising three approximately equal parts (Figure 1). A substantial head shield covers the hypostome and five pairs of segmental appendages. Seven narrow tergites, each associated with a pair of appendages, comprise the mid-trunk. This then tapers as four deeper abdominal tergites, together overlying 25 pairs of appendages**^11^**. Here we focus on the head shield, its embedded visual systems and its underlying fossilized neural components.

**Figure 1.**
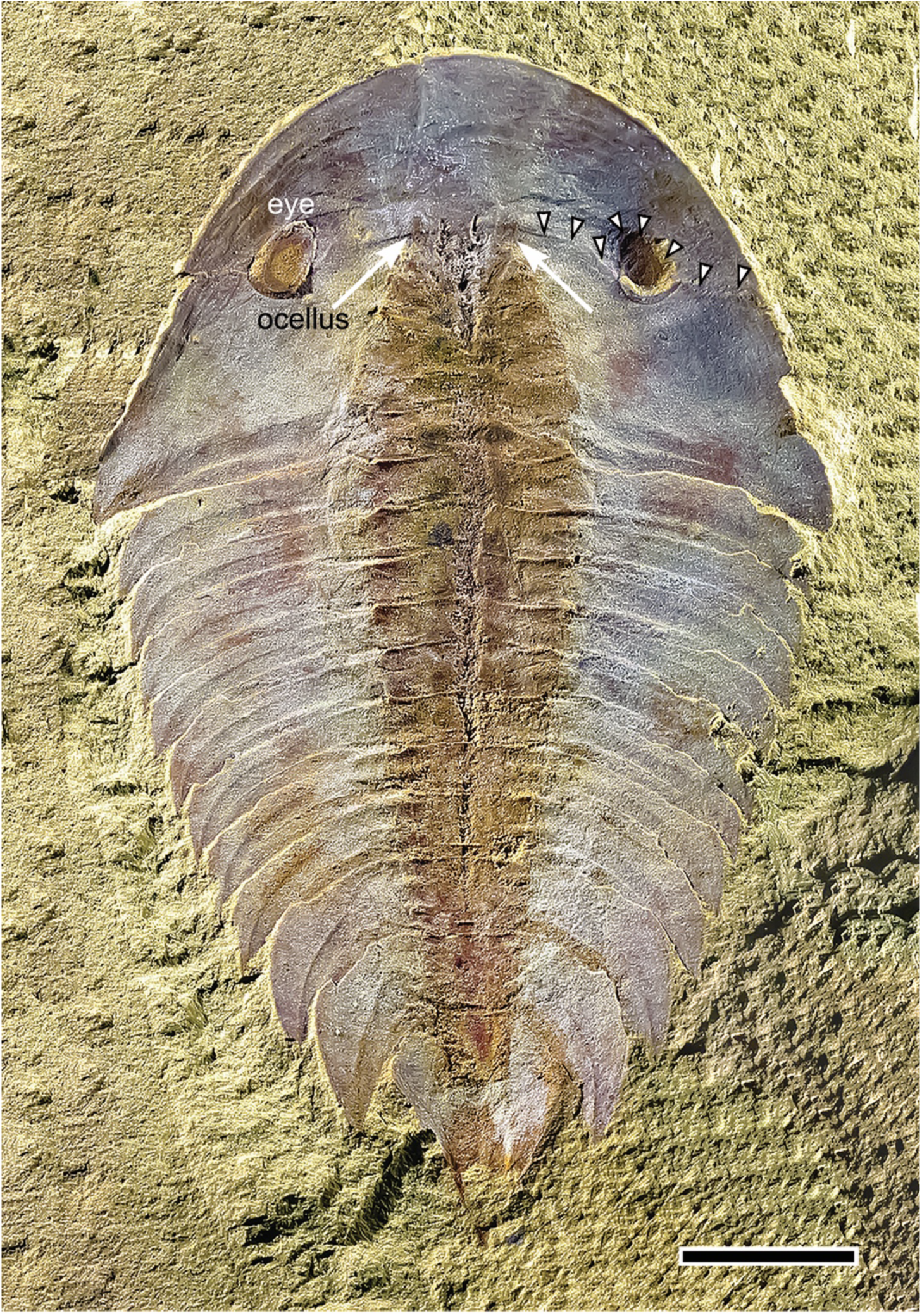
Dorsal view of *Xandarella spectaculum* (CN 115286). Prominent lateral eyes (eye) serving the protocerebrum are located at the caudal edge of the head shield’s anterior tergite. The two lateral eyes are equidistant from a medial area flanked by two small oval visual units (white arrows) interpreted as ocelli (see Fig. 2A, E, G; Suppl. Fig. S4). The seam between the anterior and the posterior tergite is indicated by arrowheads. Scale bar: 1 cm

The large head shield could be viewed as two coalesced tergites that do not obviously overlap but appear fused. The back margin of the anterior sclerite just visible in Figure 1 curves obliquely forwards to cross the midline. Its lateral passage from the midline to the edge of the shield is interrupted about half-way across by an oval socket within which resides a substantial eye (Fig. 1). The socket is not closed but abruptly narrows into a cleft extending from the eye to the lateral margin of the head shield. This is a unique attribute of this species**^14^**. The “eyestalk” is integrated ventrally within the head shield resolved as covered by a raised conduit (Fig. 2A).

**Figure 2.**
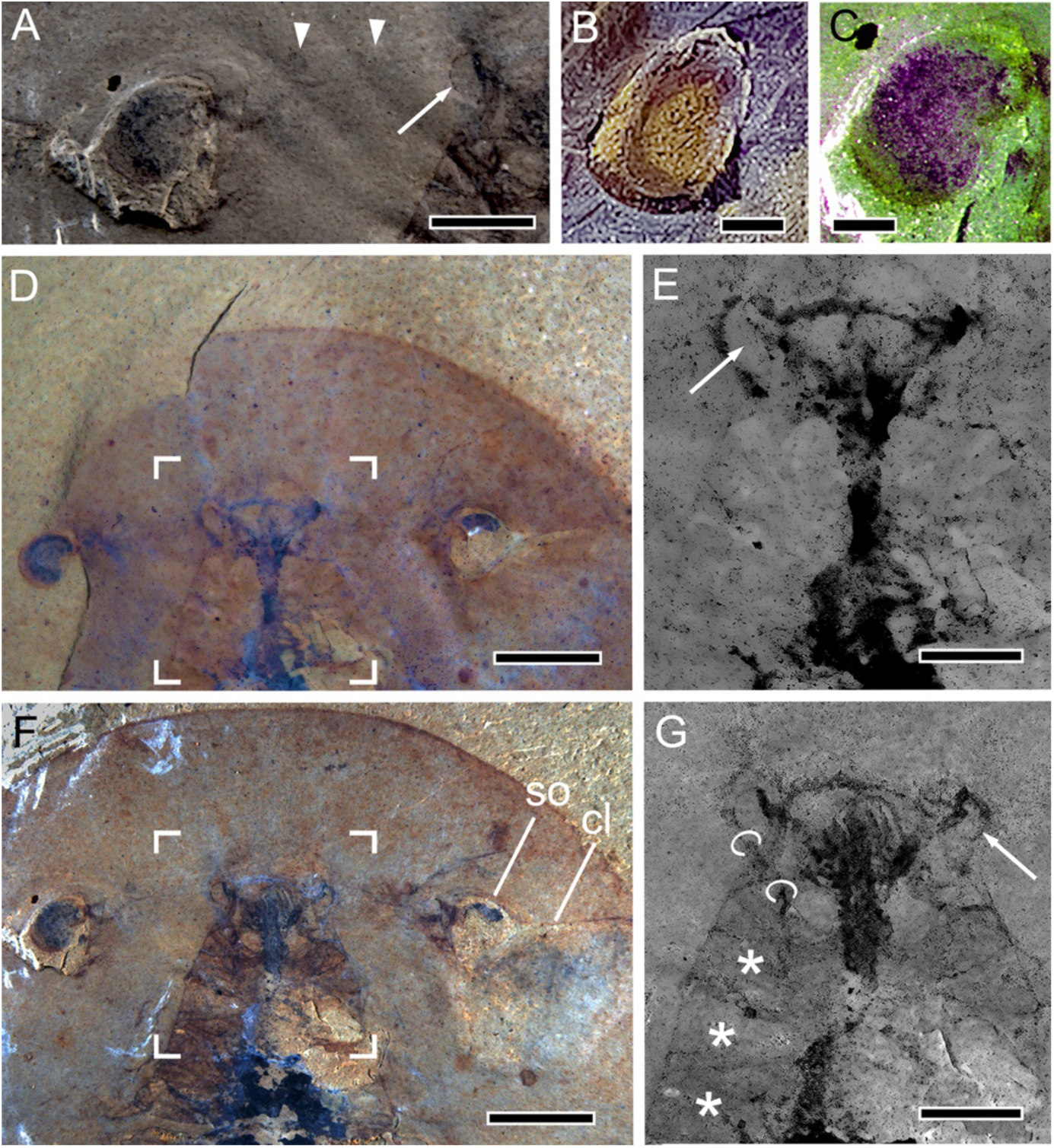
The part and counterpart of specimen YKLP 11356. **A.** The location of the optic nerve is indicated by the conduit containing the cryptic eyestalk beneath the overlying head shield; arrowheads indicate the conduit’s, hence eyestalk, trajectory. The arrow indicates the ocellus. **B, C.** In panel B, the left eye of CN 115286 shown in Fig. 1 is uncovered and shows little evidence for a compound eye; however, color substitution (panel C) of the densest part of the left eye of YKLP11356 suggests a palisade of repeated units along the eye’s upper margin. **D**. Part (rotated 180 degrees around its anterior-posterior axis to facilitate comparison with the counterpart), showing the left eye, isolated here, but visible in situ in the counterpart-. **E**. Individually separating colors as adjustable gray levels resolves details of cerebrum. The arrow indicates the left ocellus. **E.** The counterpart shows the left eye within its socket and the empty socket (so) of the right eye with the lateralized cleft (cl), as well as the cerebrum (bracketed and enlarged in G). **G.** Fossilized cerebrum with furrowed outlines of the protopod and basis (asterisks), the first three of five true segmental appendages beneath the head shield. The arrow indicates the right ocellus; rings indicate the trajectory of left antennular nerve. Scale bar F, D = 5 mm; A, E, G = 2mm; B, C = 1mm

It has been reported that the lateral eyes of *Xandarella* are nestled in bulges of the head shield and thus view ventrally**^14^**. This assumption follows observations of the related species *Sinoburius lunaris,* where downward-viewing eyes has been established**^15^**. In most fossils of *Xandarella* the lateral eyes are visible both from above or beneath, suggesting that if the eye was in life overlaid by cuticle, as has been assumed**^14^**, the cuticle must have been taphonomically fragile and easily displaced. Nevertheless, in some specimens viewed from above the eye appears nestled in a raised patch of cuticle (Fig. 2A, B, F) partially hooded by a cuticular elevation (left inset, Figure S1), both conditions partially obscuring the upper field of view. Even if the eye’s receptive layer may have been mainly directed downwards to the substrate on which the animal walked, this was probably not the eye’s exclusive panorama. Nor have observations of *Xandarella* or the related trilobitomorph *Cindarella eucala***^16^** established whether eyes claimed as comprising more than 2000 dioptric units comprise faceted ommatidia or share a transparent hemisphere with inward protruding corneal cones, as in the single-lens eyes of the Cambrian chelicerate *Alalcomenaeus* and in the lateral superposition eye of *Limulus polyphemus***^2, 17^**. Irrespective of its true field of view or resolving power, fossilized traces of the xandarellid eye show no evidence of facets nor a lamina-like peripheral neuropil that typifies extant Xiphosurans and pancrustaceans**^18, 19^**.

The lateral eyes of *X. spectaculum* are not its only visual sensors. Flanking the midline at the back margin of the anterior tergite is a pair of mirror-symmetric ocellate reliefs, indicating a pair of lensed ocelli (Figs. 1, 2A, E, G; Suppl. Figs S2, S4). These are likely homologues of what have been termed “frontal organs”**^20^**, usually paired ocelli situated on the anterior sclerite such as that belonging to the artiopodan *Helmetia expansa***^20^**. Here the ocelli are indicators of the xandarellid prosocerebral domain as they are in the prosocerebra of extant pancrustaceans**^10^**. In blind naraoiid trilobitomorphs, visible bulges anterior to the hypostome suggest cup-like accommodations of double or triple ocelli viewing upwards**^20^**; and in certain trilobites ocelli (called median eyes)**^21^** have been resolved anterior to their compound eyes. The location of ocelli in *Xandarella*, just anterior to the back margin of the anterior tectum, suggests that the latter may be a greatly expanded homologue of a discrete anterior sclerite (rostral plate) as in Helmetiidae.

## Sensory organization, brain and ventral ganglia

The part and counterpart of specimen YKLP 11356 indicating cleavage slightly tilted around the anterior-posterior axis (Figure 2D, F) reveals the fossilized neural tissue of *X*. *spectaculum*. In the counterpart the eye is nestled within a depression of the head shield where it is bracketed by cuticle at the margin of the eye’s socket (Fig. 2D, F; also Fig. S1). In the part, its more dorsal level shows the attachment of the eye to the eyestalk (Fig. 2D. Figure 2A indicates the trajectory of the channel buried in the head shield that accommodates the optic nerve. The channel is thus an eyestalk embedded within the shield leading from the eye socket to the cerebrum. Although the xandarellid eye has been suggested as comprising some hundreds of facets**^11^**, these have not been resolved with any certainty. Color substitution of the assumed receptive layer of the eye (Fig. 2C) indicates clumped arrangements as well as some marginal columnar-like elements of what might have been photoreceptors. If organized as a retinotopic mosaic these might have supported acute vision, a function already evolved in stem group Radiodonta, which are predatory arthropods. But radiodontan eyes are huge**^22^** and unlike *Xandarella*, the radiodont visual system is served not only by hexagonal arrays of thousands of facets but also a sequential arrangement of optic neuropils nested in series from immediately beneath the retina through to the midbrain**^23^** thereby demonstrating that a sophisticated object- and motion-discriminating visual system requires more than large eyes. Fossilized neuropils suggest nothing comparable in *Xandarella* as described presently.

Both the part and counterpart of YKLP 11356 support a bilaterally symmetric organization of cerebral traces resolved as dark profiles against a brown matrix (Fig. 2D, F). In Figure 2E, G these profiles are enlarged and digitally clarified by adjusting gray tones for each color range of the CMYK palette. Step focused serial images capture sharp images of all the profiles in the part and counterpart (Fig. 3A, B; Fig. S2A, B). Mirror images of each level were used to reconstruct the bilaterally symmetric cerebrum, detailed in Figure 3C and Suppl. Fig. S2A, B. The brain is clearly organized as three discrete levels of elaboration along its anterior-posterior axis. Most anteriorly, a neural arch interpreted as the prosocerebral bridge links laterally situated apical lobes and sends several delicate and slender nerves downwards into the underlying neuropil, interpreted as the flattened medial volume of the protocerebrum (Suppl. Fig. S2B). The two apical lobes are associated with black granular deposits that outline the afore-mentioned lenticular reliefs interpreted as ocelli (Fig. 2E, G). The protocerebrum is further defined by a pair of diminutive optic lobes that are supplied by the optic nerve emerging from its conduit (Suppl. Fig. 2B, C). The protocerebral domain is further defined by a pair of recurrent columnar lobes extending forward toward the prosocerebrum. Because the brain is essentially flattened it is uncertain whether the lobes merge with the prosocerebrum. Further caudally, and mainly resolved in the counterpart, a cluster of glomerular traces resolves a paired ensemble of large, flattened glomeruli from which originate the paired antennular nerves (panels *cp* Fig. 2A; lower two panels Suppl. Fig. 2B, D, E). supplying flagelliform antennules. These with their glomerular origin define chemosensory–haptic neuropils that in extant pancrustaceans (Hexapoda+Crustacea) are constrained to the brain’s deutocerebrum**^10^**.

**Figure 3.**
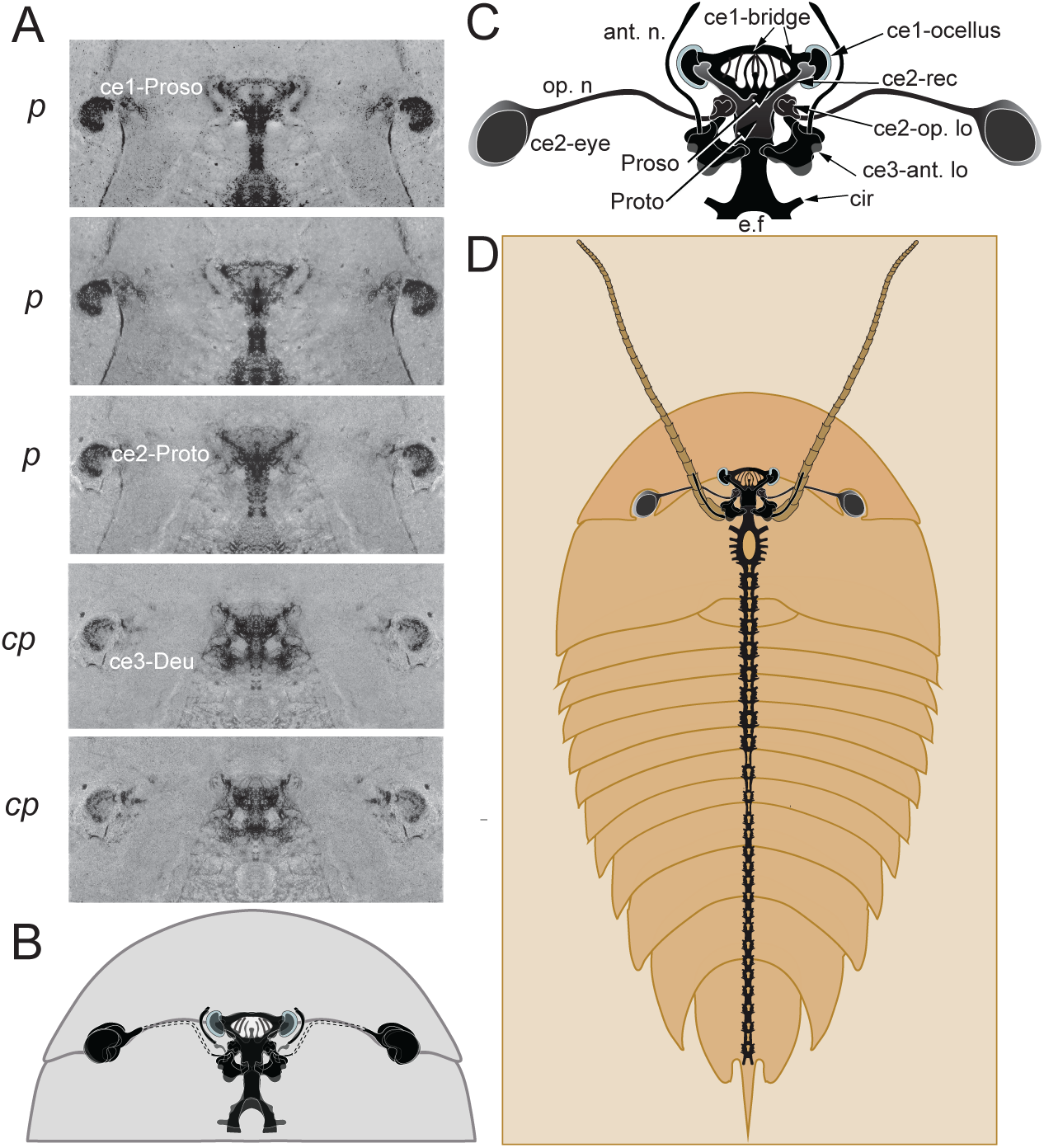
Xandarellid cerebrum and ventral ganglia. **A**. Serial photographs of specimen YKLP 11356 shown as mirror-symmetric profiles (see Suppl. Fig. S2). The panels show stepped focal images of fossil tracts and neuropil at three levels of the part (p) and at two levels of the counterpart (cp). **B**. Reconstruction obtained from overlapping silhouette-type infill of each image (for details see Suppl. Fig. S2). The double dashed line indicates the optic nerve conduit. **C**. Enlarge reconstruction: neural traces resolve a pair of lobes beneath the paired ocelli connected by a bridge that sends slender connections to the underlying medial prosocerebrum. Optic nerves leading from the lateral eyes can be extrapolated from their head-shield conduit. Optic nerve end at a diminutive terminal lobe in protocerebrum. These are dwarfed by the paired antennular lobes at the level of the deutocerebrum (ce3). The protocerebrum also provides a pair of recurrent lobes that partially overlap ce1. The caudal extension of the nerve appears smudged, but its initial trajectory indicates that the ensuing ventral cord splits into two tracts. **D**. The tracts are shown encircling the esophageal foramen and then extend ventrally providing segmental ganglia typifying the sister trilobitomorph species *Cindarella eucalla* shown in Figure 4. Abbreviations: ce1-bridge, prosocerebral bridge (domain ce1); ce1-ocellus, one of the paired prosocerebral ocelli; Proso, rostral prosocerebrum (ce1); medial Protocerebrum (ce2); ce2 eye, protocerebral eye; op. n, reconstructed optic nerve; ce2–op. lo., protocerebral optic lobe; ce2-rec, recurrent protocerebral lobes; ce3–ant. lo, deutocerebral antennular lobe; ant. n., antennular nerve; e. f, esophageal foramen; cir, initial part of the circumesophageal nerve.

Specimen YKLP 11356 provides part of its caudal extension towards the ventral nerve cord, which splits into two halves to extend around the gut. The reconstruction of a complete ventral ganglionic central nervous system shown in Figure 3D includes arrangements from the trilobitomorph *Cindarella eucalla* (YKLP 11141) shown in Figure 4. *Cindarella eucalla* is the sister taxon of *X. spectaculum*; it has well-defined segmental ganglia (center *B’* in Fig. 4A; B-D) which are here ‘adopted’ to reconstruct a complete trilobitomorph central nervous system (Fig. 3D). YKLP 11141 also shows a partially preserved head shield bordered by a laterally positioned eye comprising what may suggest aligned visual units (Fig. 4A, panel *A’*), as suggested in an earlier description of a different specimen of *C. eucalla***^20^**.

**Figure 4.**
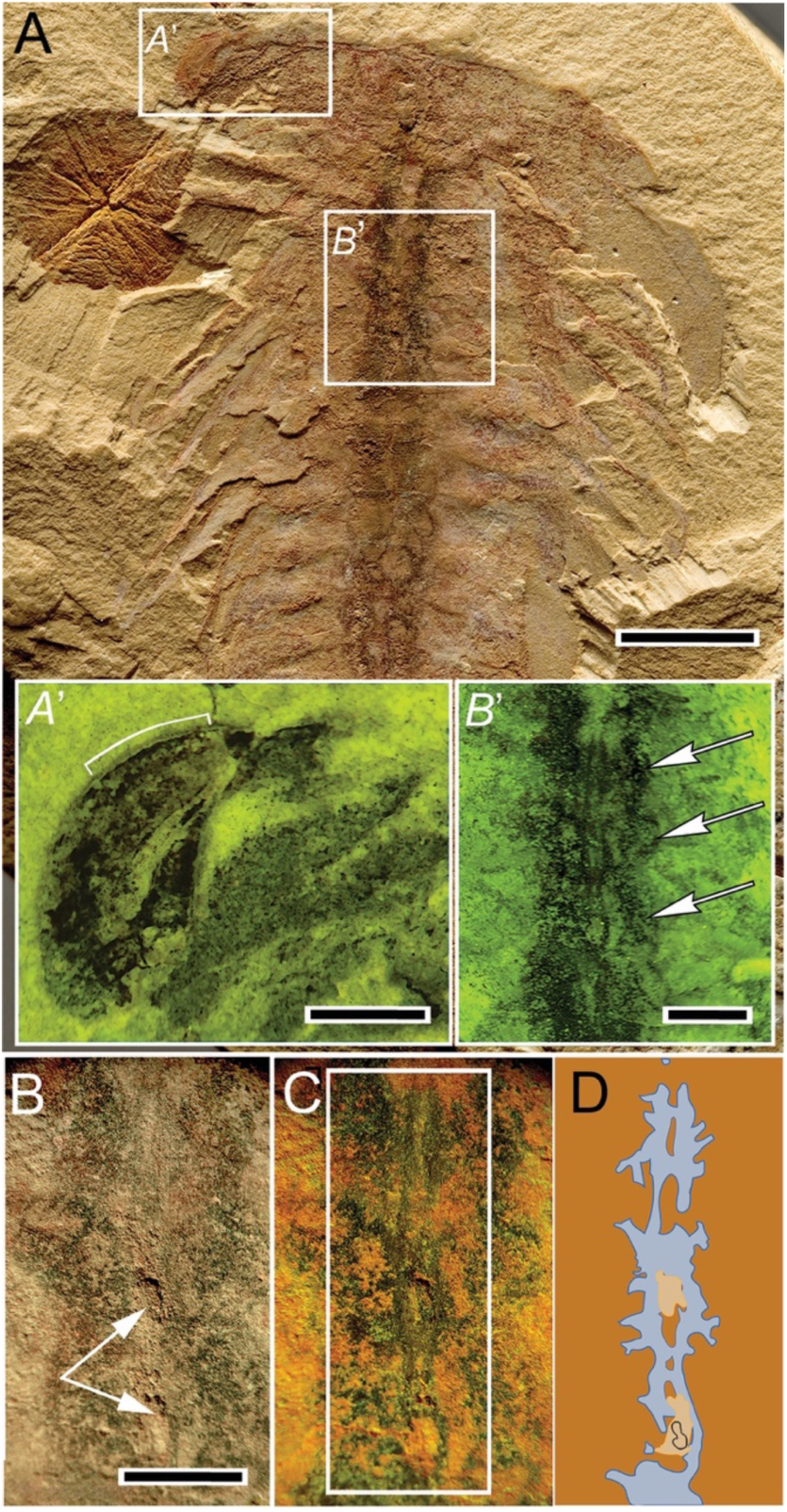
*Cindarella eucalla* segmental ganglia. **A.** Top-down view of YKLP 11141 showing discrete segmental ganglia underlying the gut. The head shield is partially preserved. A lateral eye (inset *A*’) flanks the head shield. The gut is indicated by segmentally arranged diverticuli and glands. **Inset *A*’.** Enlargement of ultraviolet (UV)-illuminated eye with serial units (bracketed) of what may be non-facetted photoreceptors. **Inset *B*’.** UV illumination reveals segmentally paired digestive glands (arrows) and caeca beneath a ladder-like arrangement of ganglia and their connectives. **B.** White light illumination suggests gut sediments have fallen out (arrowed) exposing the gut’s approximate width. **C.** Combined white light and UV resolve the gut and the ventral chain of ganglia. **D**. Tracing of ganglia from boxed area in panel C. Their diagrammatic representation has been ‘married’ to the cerebral nervous system of *Xandarella* shown in Fig. 3D. Scale: A = 5mm, *A*’=1mm; *B’* = 2mm; B, C, D = 1mm.

## Comparative alignments: empirical evidence of evolved loss

Fossil cerebra can be aligned with cerebra of extant arthropods using as common reference points the endomesodermal interface situated between the deutocerebrum and the first hox-defined true segment variously called the tritocerebrum or intercalary segment**^24^** (Fig. 5). Sensory attributes denoting each of the three cerebral domains (ce1, ocelli; ce2, lateral eyes; ce3, paired deutocerebral appendages; Fig. 5B) provide further information by aligning cerebral domains and their defining neuropils with combinations of gene expression that determine them (Fig. 5A); developmental genetic studies of mandibulate arthropods demonstrate that specific computational neuropils characterize each cerebral domain**^10, 24–26^**. Their volume and elaboration reflect their prominence in mediating specific behaviors as shown in numerous studies exemplified by five here**^27–31^**. The prosocerebrum (ce1), which provides the ocelli and incorporates the central complex is prominent in those taxa that employ allocentric exploration and homing using path integration**^27, 28^**. The protocerebrum (ce2), which is served by the lateral eyes, is also defined by its cognate optic neuropils, multisensory centers, and the paired mushroom bodies which support learning and memory**^29–32^**. The comparative size and diversification of centers defining each domain reflect their roles in vision and memory-dependent behaviors. Likewise, degrees of elaboration of the antennular lobes of the ce3 domain (the deutocerebrum) reflect their importance in chemosensory-driven behaviors**^33,34^**.

**Figure 5.**
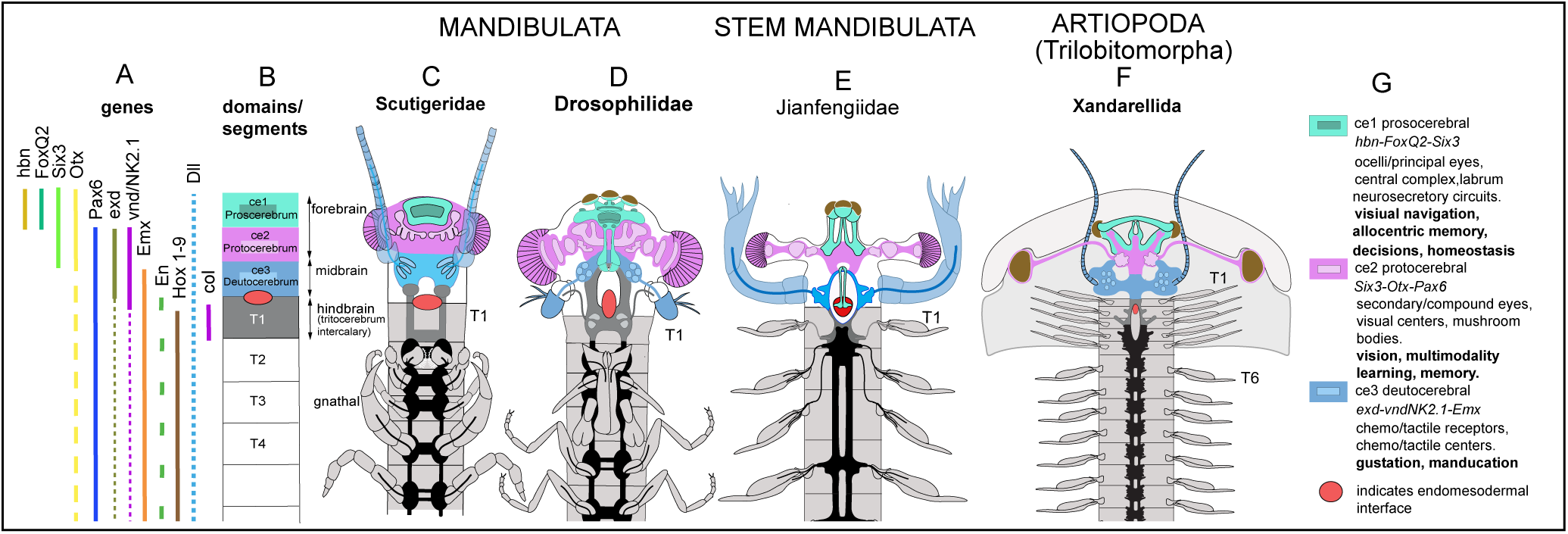
Alignment of extant mandibulate, stem mandibulate and trilobitomorph brains. (**A**) In living euarthropods, the combinatorial expression of homologous genes (for source references see Strausfeld et al.**^24^**) defines three cerebral domains (**B**): these are the prosocerebrum (ce1); the protocerebrum (ce2); and the deutocerebrum (ce3). Specified by the gene *collier***^44^**, T1 is the first true segment of the body. Notably, the cerebral traits defining each lineage are independent of Hox-code determination of trunk segmental tagma such as T2-T4, which contribute gnathal appendages in extant mandibulates. (**C-F**) Schematic showing the morphogenetic divisions of the cerebra of two extant mandibulates (**C, D**) aligned with the corresponding neuroanatomical domains of the stem mandibulate *Jianfengia multisegmentalis***^9^**(**E**) and the trilobitomorph *Xandarella* (**F**). The linear brains of both Cambrian and extant mandibulates are characterized by their sensory systems and computational centers (summarized in **G**). The trilobitomorph cerebrum is unusual in that its protocerebrum is exceptionally small. Notably, unlike crown and stem mandibulates (**D, E**) no neuropils underlie the trilobitomorph retinas, whose optic nerves supply simple visual centers suggesting a single neuropil. An extension from the protocerebrum forwards to the prominent prosocerebrum indicates that simple visual cues would converge with signals encoded by the paired ocelli. These arrangements together with the prosocerebral bridge suggest that ocular and ocellar neuropils contributed to navigation entailing path integration**^38,39^**, as occurs in extant crustaceans. The paired antennules of *Xandarella* supply information to prominent glomerular deutocerebral lobes larger than those of extant pancrustaceans, taking into account taphonomic flattening. Matches between exceptional fossilized brain regions and corresponding neuropil enlargements in extant pancrustaceans are shown in Suppl. Fig. S3.

When aligned against extant and coeval taxa (Fig. 5C-F), the xandarellid brain is shown to generally conform to the expected euarthropod cerebral ground pattern of three linear domains defining the proso-, proto-, and deutocerebrum**^10, 24^**. But the xandarellid cerebrum is nevertheless strikingly different in that its second domain, the protocerebrum, is exceptionally small (Fig. 5F). Notably, unlike crown and stem mandibulates no neuropils have been found that underlie the lateral eye retinas whose optic nerves each supplies a diminutive visual center indicating a single synaptic neuropil. Extant examples arrangements where large eyes serve simplified visual systems are provided by scutigerid centipedes where prominent compound eyes send connections to a small single layered centers centrally the organization of which suggests that likely responds to the velocity but not direction of visual flow**^35^** (Figure. 6).

**Figure 6.**
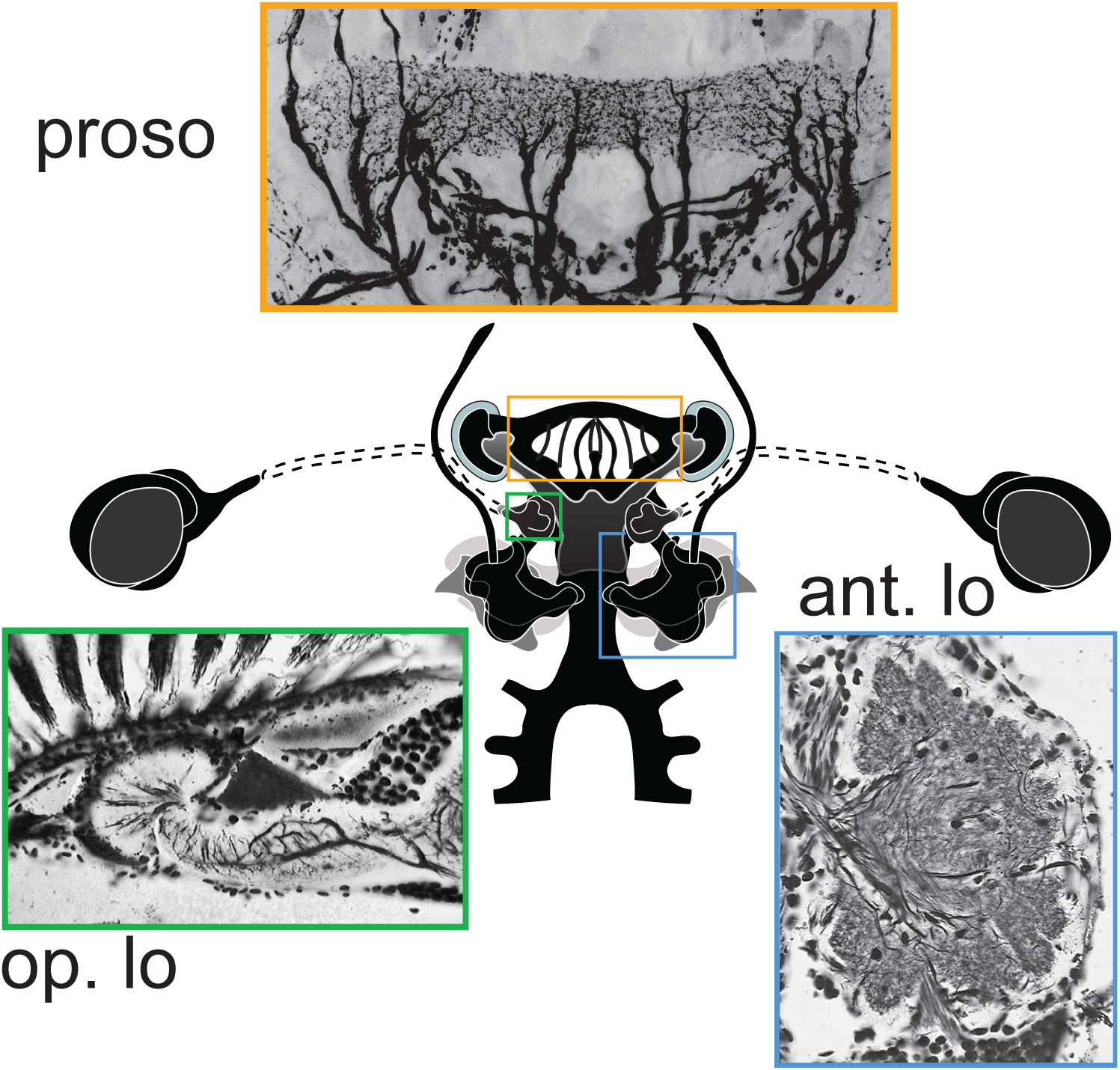
**Prominent silhouette-like depositions signifying fossilized neuropils in *Xandarella spectaculum* that correspond to comparable centers in mandibulates (clockwise from the top). proso**: The neural traces of xandarellid prosocerebral arch correspond to the orderly arrangement of relay neurons in the pancrustacean bridge that extend downward axons into the underlying neuropil, here show from a larval odonatan. **ant lo**: The voluminous glomerular lobes of *Xandarella* corresponds to the antennular lobes of stomatopods, benthic predators renowned for their homing behaviors. **op. lo**: Despite its large lateral eyes, *Xandarella* has diminutive optic neuropils. An extant example of such disparity is provided by scutigeromoph centipedes which despite possessing substantial eyes have small unistratified optic lobe neuropil.

## DISCUSSION

Despite the absence of reliable information about their brains the same cerebral ground pattern shown in Fig. 5 is indicated for Trilobita. The possession medial ocelli**^21^** in some trilobite species as well as having lateral compound retinas and paired antennules indicate the presence of three cerebral domains. Further, trilobite morphology is so similar to other trilobitomorph soft–bodied species that the xandarellid brain can be taken, at present, as typifying the entire subphylum. An extension from the protocerebrum forwards to the prominent prosocerebrum (Figs. 3C, 6; Suppl. Figs. S2E) indicates that visual information such as visual flow was integrated with polarized light signals encoded by the paired ocelli, as occurs in extant pancrustaceans**^36, 37^**. The xandarellid ocelli are partially enwrapped by the prosocerebrum’s lateral lobes (Fig. 2E, G) which are linked heterolaterally by the arch-like neuropil connected to the underlying protocerebrum by a row of slender tracts (Figure 3A, C, 6; Suppl. Figs. S2). This discrete organization immediately indicates homology with the ordered structure of the pancrustacean prosocerebral bridge (Fig. 6) which in extant pancrustaceans is a crucial component that computes and records allocentric directional headings, thus determining egocentric path integration**^38^**. The recognition of an evolved diminution of the protocerebrum coupled with an exaggerated prosocerebral bridge and huge chemosensory lobes, also typifying benthic crustaceans (Suppl. Figs. S2, S3), allows considerations about what kind of behavior was supported by the xandarellid brain and whether its cerebral organization could resist ecological stress.

The expanded dimension of the xandarellid prosocerebral domain is telling. In modern pancrustaceans the elaboration of the prosocerebral bridge reflects their expanded role in multimodal integration in foraging and navigation**^38,39^**. Today’s marine stomatopods and isopods, many of which are benthic predators or foragers, possess the most elaborate central complexes where an archlike bridge maps a series of axon bundles into the underlying central body**^28^**. Here in *Xandarella* the neural traces are so well-preserved as to indicate a strikingly similar arrangement of regular channels between the bridge and underlying prosocerebrum (Fig. 6). As remarked above, the prominent and detailed prosocerebral bridge of *Xandarella* starkly contrasts with its much smaller protocerebrum. This is an important distinction from extant pancrustaceans whose protocerebrum is voluminous not only because of its sequential visual neuropils but also because of its other sensory circuits including, in particular, its paired mushroom body memory centers**^10,31,32^**. The sparse protocerebrum of *Xandarella* suggests simplicity; meagre optic neuropils with few other attributes other than recurrent lobes extending forwards to the prosocerebrum (Fig. 2A, C), reminiscent of exiguous mushroom bodies as seen in extant Collembola**^40^**.

The xandarellid cerebrum, despite sharing the ground pattern of three defined domains is clearly set apart from other fossilized mandibulate brains**^1,6,9^**. Its conspicuous olfactory/chemosensory lobes and its elongated antennules for assessing benthic chemical and tactile cues denote that odorant and haptic discrimination played the major role in trilobitomorph behavior and survival. Abundant studies of pancrustacean behavior has shown that foraging also requires information about space and its limitations. As described from observations of extant pancrustaceans, paired ocelli and lateral eyes together mediate navigation; even at low light intensities changes in orientation of polarized light can function as compass-like references informing the brain about rotational change of direction**^37^** as can sun compass orientation guide foraging isopods**^41^**. Integrated with information from the eyes that compare left and right visual flow velocity provide further information about path integration and thus movements within a circumscribed ecology suited to the species. Evidence from soft tissue remnants in various trilobitomorphs**^42^** showing gut contents is suggestive of benthic scavenging. Yet the overall shapes of trilobitomorphs and their evolutionarily reduced cerebral attributes also suggests that they were poorly equipped to adapt to climatic and thus ecological change. *Xandarella* went extinct before the Ordovician. The final demise of trilobites was delayed until the major ecological disruptions defining the end of the Permian**^43^**.

## METHODS

### Material Provenance

Descriptions herein refer to specimens retrieved from the Cambrian (Series 2, Stage 3) Eoredlichia-Wutingaspis trilobite biozone, Yu’anshan Member, Chiungchussu Formation, Haikou. The specimens used here are curated at the Yunnan Key Laboratory for Palaeobiology (YKLP), Institute of Paleontology, Yunnan University, Yunnan, Kunming, China.

### Photomicroscopy

For light microscopy of fossil material, digital images were taken using a Nikon D3X attached to a Leica M205C photomicroscope (Leica Microsystems; Wetzlar, Germany). Images were transferred to Adobe Photoshop CS5 (Adobe Systems; San Jose, CA) and processed using the Photoshop camera raw filter plug-in to adjust sharpness, luminance, texture, and clarity. Unless otherwise stated, colors were untouched and are those typical of Chengjiang fossils. For reconstructing neural traces serial images focusing at micron-intervals capture all the relevant profiles in both part and counterpart of specimen YKLP 11356 as shown in Fig. 3A and in supplemental Figure S2. These were used to reconstruct the bilaterally symmetric cerebrum, detailed in Figure 3B. For ultraviolet illumination (Figure 4A’,B’,C), fossils were photographed using appropriate filter blocks to evoke intense green fluorescence. Flexible fiberglass light guides were used to combine UV fluorescence and white-light illumination.

### Isolating neural traces in specimen YKLP 11367

Cerebral traces initially resolved as dark profiles against an intensely dense brown background were digitally clarified by converting color images to monochrome, using the Adobe Photoshop Black & White function for luminosity adjustments of each color of the CMYK palette.

### Interpretive drawings and tracings

Drawings were made using Adobe Illustrator for which layered tracings and reconstruction were generated using photographic images as guides.

## Data Availability

Specimens used or referred to here were curated by X. H. at the Yunnan Key Laboratory for Palaeobiology (YKLP), Institute of Paleontology, Yunnan University, Yunnan, Kunming, China. Data reported in this study are included in the article and the supplementary material. Any additional information regarding the data reported in this article is available from N.J.S. upon reasonable request.

## Code Availability

This article does not contain original code.

## ACKNOWLEDGEMENTS

Our special thanks go to Dr. Camilla Strausfeld for crucial advice and critical editing of the manuscript. We have profited from Dr. Greg Edgecombe, London Museum of Natural History, for his suggested emendations, from Dr. Lorenzo Lustri, Universities of Lausanne, Switzerland and Kunming, China for his critical reading and advice, and from Dr. Bruce Lieberman Department of Ecology & Evolutionary Biology, University of Kansas, for his reading the work. We thank Professor Eric Warrant, Department of Biology University of Lund, Sweden, for his valuable advice regarding the possible visual receptivity and behavioral range afforded by the xandarellid visual system. Marc Haensel B. A., of Paleoarchives, Montreal, Canada generously provided an image of *Xandarella* used here for Suppl. Fig. S1.

## FUNDING

Funding for this work was enabled by residual funds from a 1998 John D. and Catherine T. MacArthur Foundation Fellowship awarded to N.J.S. F.H. acknowledges support from the UK Biotechnology and Biological Sciences Research Council (BB/N001230/1).

## AUTHOR CONTRIBUTIONS STATEMENT

N.J.S. and F.H. originated the project; X.H. discovered and named the species, provided specimens and with N.J.S. discussed the material. N.J.S examined and photographed the fossils YKLP 11356, YKLP 11141. F.H. ascribed published gene expression data to the described panarthropods. N.J.S. and F.H. wrote the manuscript with input from X.H.

## CONTACT AND COMPETING INTEREST INFORMATION FOR ALL AUTHORS

The authors declare no competing interests.

## SUPPLEMENTAL INFORMATION

**Figure S1.**
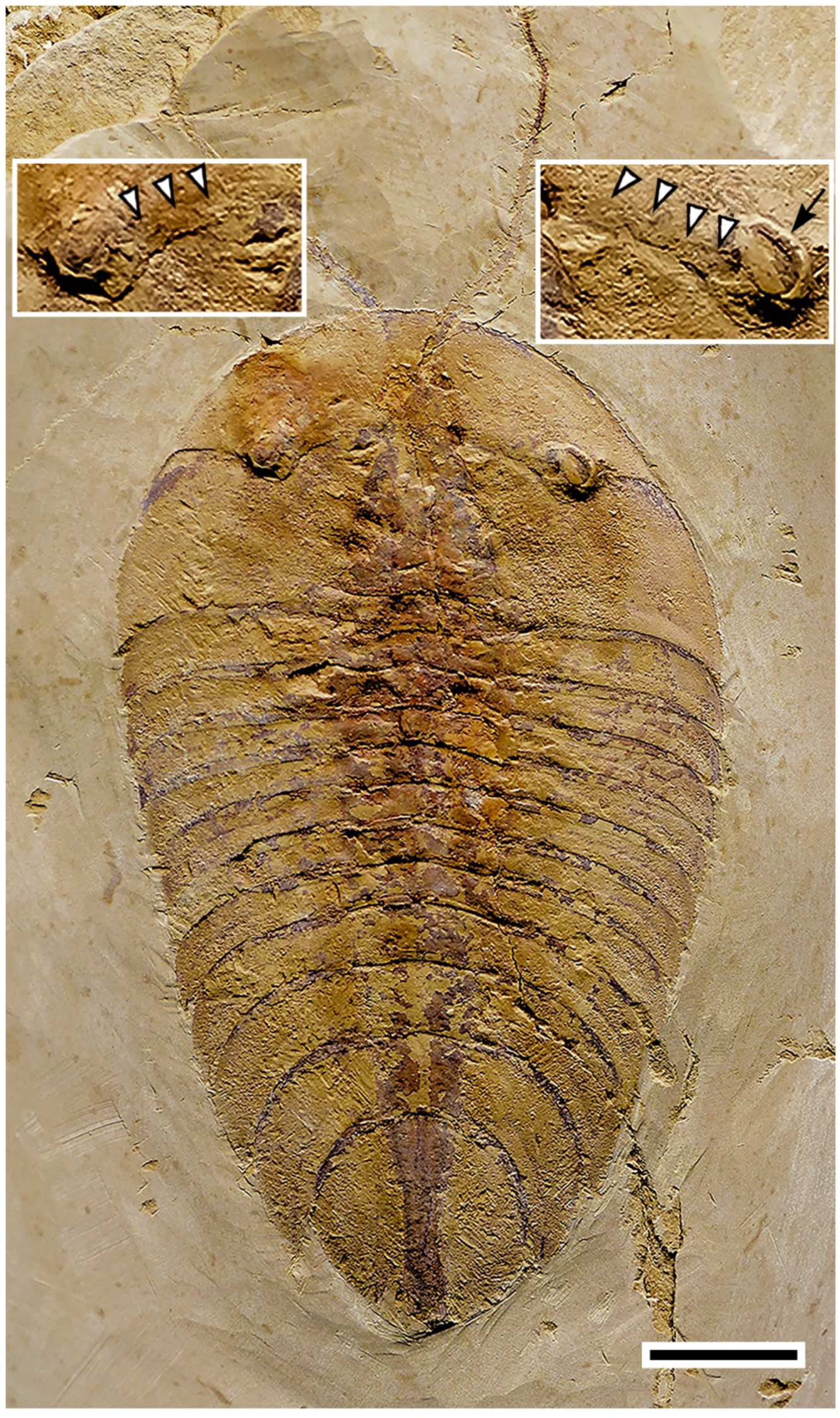
Dorsal view of *Xandarella spectaculum*. The insets show the lateral eyes completely located in the head shield cuticle. Both sides show the slightly raised conduits (arrow heads in insets) that house the embedded optic nerve. The right side indicates the laterally directed surface of the (arrow) extending above the level of shield. Scale bar = 1 cm. (see Acknowledgements).

**Figure S2.**
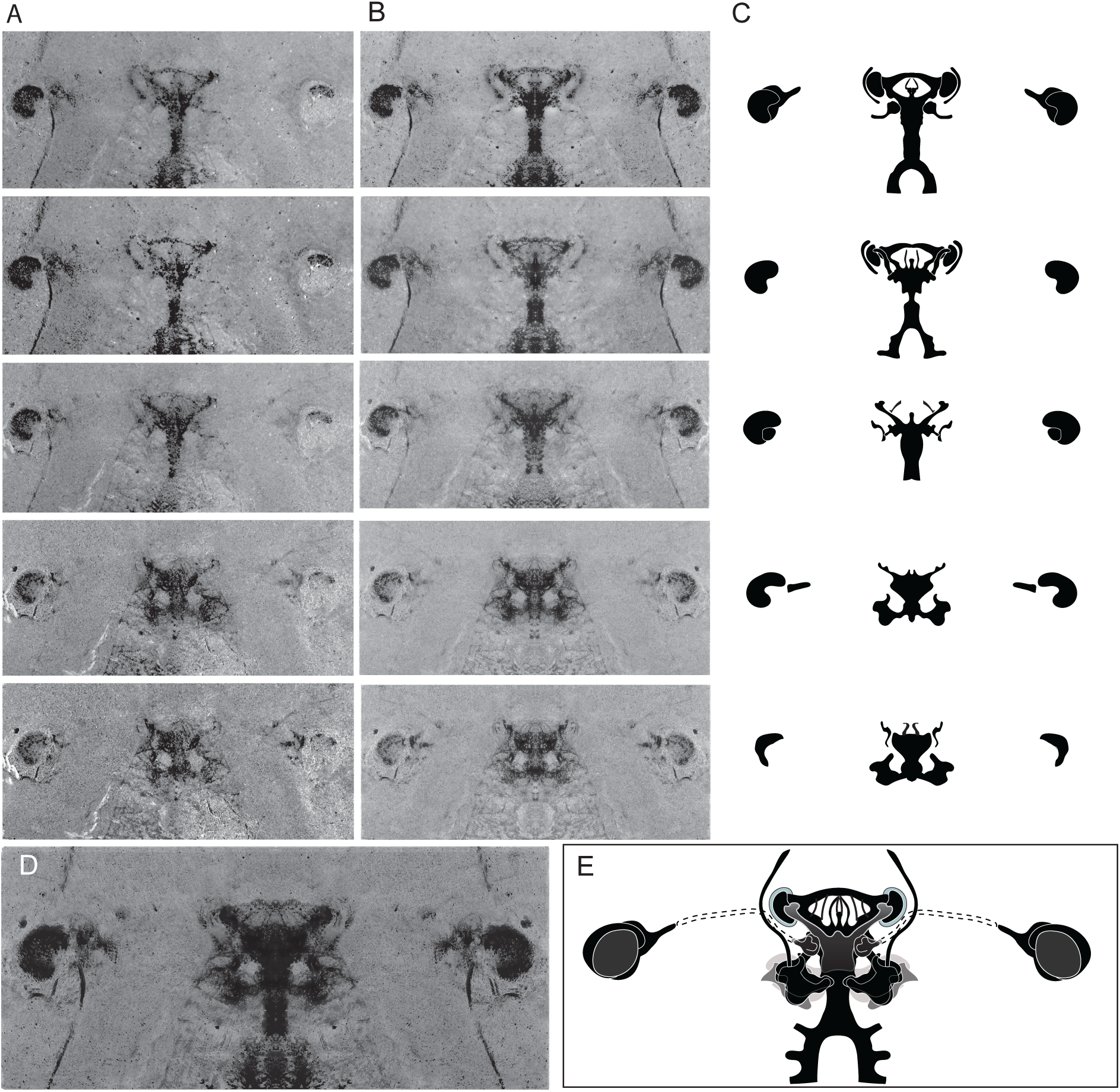
Reconstruction of the Xandarellid brain (YKLP 11356). **A**. Serial photographs of the part (upper 3 frames) and counterpart (lower 2 frames). **B**. Each frame in A is copied, flipped horizontally, and superimposed over itself to provide a mirror symmetric image. **C**. Silhouette-type infill of each image in B. **D**. Guideline image of all frames in B superimposed. **E**. Final reconstruction. Cryptic optic nerve trajectory within the overlying head shield is indicated as dashed lines.

**Figure S3.**
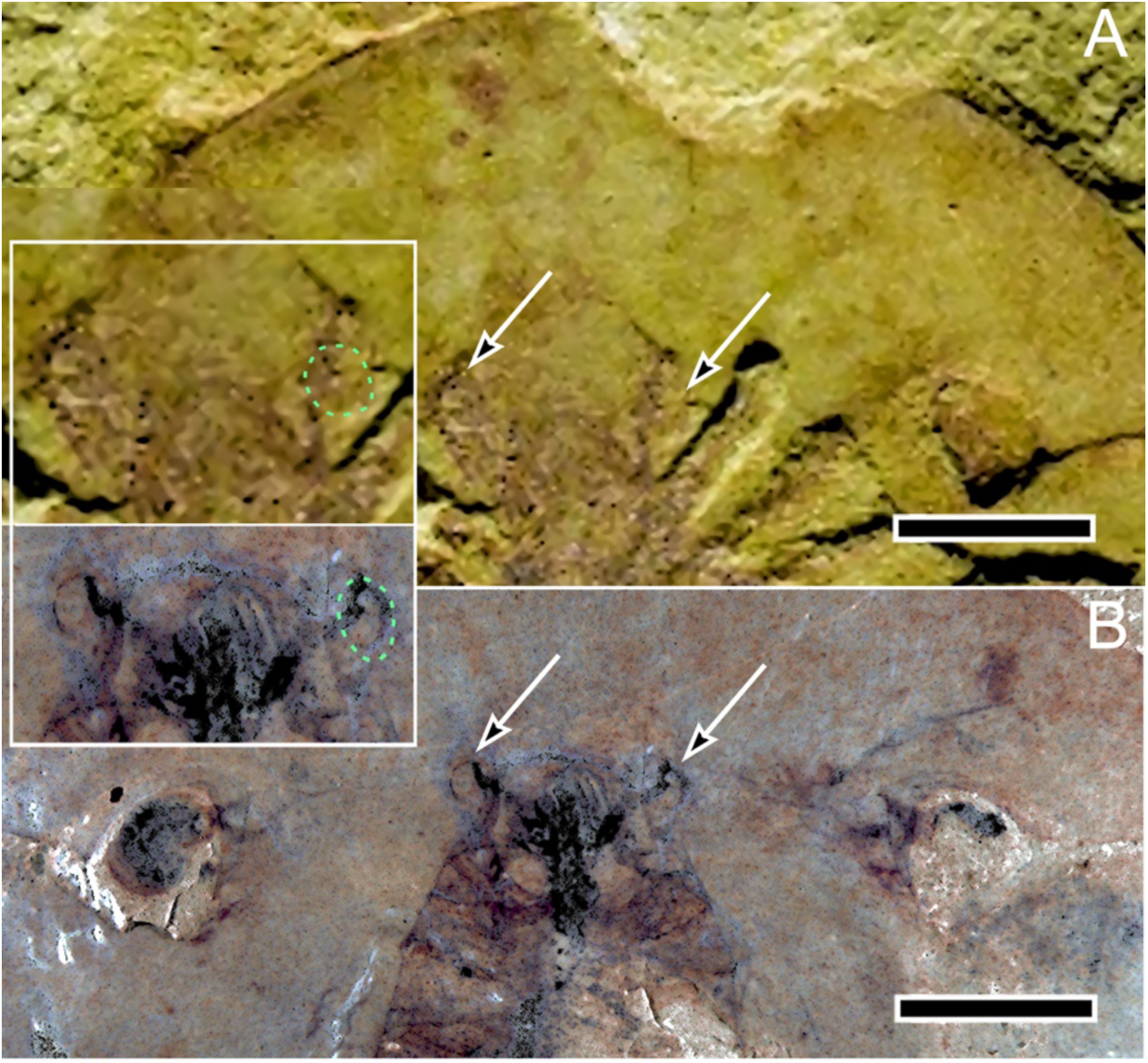
*Xandarella spectaculum* ocelli. A (NIGPAS 115286), B (YKLP 11356). Two specimens of *Xandarella* demonstrate the paired ocelli (arrowed) at the back edge of the head shield’s anterior tergite. The right ocellus in each enlargements is outlined. Scale bars A, B: 1 cm.

## Notes

### Competing Interest Statement

The authors have declared no competing interest.

### Summary of Updates

Reorganized text to align with reorganized and expanded number of figures and their legends in the main narrative shifted from the now reduced supplemental information.

